# Inhibitor of sarco/endoplasmic reticulum calcium-ATPase impairs paramyxovirus replication

**DOI:** 10.1101/328997

**Authors:** Naveen Kumar, Sanjay Barua, Nitin Khandelwal, Kundan Kumar Chaubey, Ram Kumar, Krishan Dutt Rawat, Yogesh Chander, Thachamvally Riyesh, Shoor Vir Singh, Bhupendra Nath Tripathi

**Author notes:** Equal contribution (Naveen Kumar, Sanjay Barua). Address correspondence to Naveen Kumar,; Bhupendra N. Tripathi,.

## Abstract

Sarco/endoplasmic reticulum calcium-ATPase (SERCA) is a membrane bound cytosolic enzyme that is known to regulate the uptake of calcium into the sarco/endoplasmic reticulum. Herein, we demonstrate for the first time that SERCA can also regulate paramyxovirus [Peste des petits ruminants virus (PPRV) and Newcastle disease virus (NDV)] replication. Treatment of Vero cells with SERCA specific inhibitor (Thapsigargin) at a concentration that is nontoxic to the cells significantly reduced virus replication. Conversely, overexpression of SERCA rescued the inhibitory effect of Thapsigargin on virus replication. PPRV/NDV infection induced SERCA expression in Vero cells which could be blocked by Thapsigargin. With the help of time-of-addition and virus step-specific assays, it was observed that Thapsigargin specifically inhibits viral entry and subcellular localization of the viral proteins. Furthermore, NDV, but not PPRV acquired a significant resistance to Thapsigargin on long-term passage (P=70) in Vero cells. To the best of our knowledge, this is the first report describing virus supportive role of SERCA and a rare report suggesting that viruses may acquire resistance even in the presence of an inhibitor that targets a cellular factor. This study will contribute in understanding paramyxovirus replication and development of antiviral therapeutics using SERCA (host factor) as a candidate drug target.

## INTRODUCTION

The control strategies against pathogens have classically relied upon targeting essential proteins of the pathogens (1). High mutation rate in viral genome allows the virus to become resistant to antiviral drugs and preexisting immunity (2). Classically, antiviral drugs have been developed by directly targeting viral proteins (3). Due to high mutation rates, virus gets mutations at the druggable targets and becomes resistant. The rise in incidence of drug resistance has prompted a shift in the development of novel antiviral drugs (4). Viruses are obligate intracellular parasites that are highly dependent on host. Host responses are equally important in determining actual outcome of the diseases. Upon viral infection, numerous cellular factors are dysregulated (increased or decreased expression); some of these host factors facilitate virus replication (proviral), whereas others may have antiviral function (5). Proviral host factors may serve as targets for development of novel antiviral therapeutics (1, 6, 7).

Protein phosphorylation and dephosphorylation, mediated respectively via kinase and phosphatases, is a ubiquitous cellular regulatory mechanism during signal transduction which determines key cellular processes such as growth, development, transcription, metabolism, apoptosis, immune response, and cell differentiation (8). Kinome-the protein kinase complement of the human genome, completed in 2002, identified 518 protein kinase genes. These kinases have been shown to play a key role in cancer and many other diseases (8) including viral infections (9), making these proteins very desirable drug targets.

In vertebrates, there are three families of P-type Ca^2+^-ATPases that regulate homeostasis of intracellular Ca^2+^ level. Plasma membrane Ca^2+^-ATPase (PMCA), SERCA and secretory pathway calcium ATPAse (SPCA) are located in plasma membrane, endoplasmic reticulum and Golgi apparatus respectively (10). SERCA transports Ca^2+^ from cytosol to the double membrane-bound (endoplasmic reticulum) intracellular compartments (11-13). SERCA is also involved in other cellular functions such as signal transduction, apoptosis, exocytosis (14), cell motility (15) and transcription (16). There are three genes (ATP2A1-3) in vertebrates that code for three SERCA isoforms, namely SERCA1-3 (17, 18). Each of these genes undergoes alternative splicing and hence results in 10 SERCA proteins (two each from SERCA1 and 2 and six from SERCA3) (19). While some of these isoforms/variants are ubiquitously expressed in most cell types (SERCA2), others show a range of cell type specific expression patterns (12, 17, 20). The role of these Ca^2+^-ATPases in virus replication is only beginning to be appreciated. Whereas the role of SERCA and PMCA in virus replication remains unknown, a recent study suggests that SPCA1 supports virus replication (21).

Previously we screened a library of kinase and phosphate inhibitors for their antiviral potential against influenza A viruses (6). Herein, we also screened library of these chemical inhibitors for their antiviral effects against paramyxovirus-morbillivirus [(peste des petits ruminants virus (PPRV)] and avulavirus [(Newcastle disease virus (NDV)]. Inhibitor of sarco/endoplasmic reticulum calcium-ATPase (SERCA) was identified as one of the candidate that blocked NDV and PPRV replication. In this study, we provide mechanistic insights into the potential roles of SERCA in facilitating paramyxovirus replication, primarily in mediating viral entry and localization of viral proteins, thus revealing SERCA as a potential target for development of antiviral therapeutics.

## MATERIALS AND METHODS

### Cells and viruses

Vero (African green-monkey kidney), 293T (human embryonic kidney), HeLa and goat kidney cells were grown in Dulbecco’s Modified Eagle’s Medium (DMEM) supplemented with antibiotics and 10% heat-inactivated fetal bovine serum. PPRV and NDV were available in our laboratory and have been described elsewhere (22). Viral titers were determined by plaque assay, as previously described (22).

### Inhibitor

Thapsigargin (SERCA inhibitor) was procured from Sigma (Catalog Number-T9033, Steinheim, Germany). Thapsigargin is a non-competitive inhibitor of SERCA. It is extracted from a plant *Thapsia garganica* and structurally classified as a sesquiterpene lactone (23).

### Antibodies

SERCA2 ATPase Antibody (MA3-919) was procured from Invitrogen, (Carlsbad, USA). Anti-PPRV and anti-NDV serum is described elsewhere by our group (24). Secondary fluorescein isothiocyanate-conjugated anti-rabbit antibody and secondary Rhodamine-conjugated anti-mouse antibody were purchased from Sigma (Steinheim, Germany).

### MTT assay

Cytooxicity of Thapsigargin was analyzed in MTT assay, as previously described (24).

### Antiviral efficacy

Vero/MDBK cells were infected with respective viruses at MOI=0.1 in the presence of 0.5 µM Thapsigargin or vehicle control (DMSO). Progeny virus particles released in the supernatants were quantified by plaque assay.

### Virucidal activity

Virus suspensions (PPRV/NDV) containing ∼10^6^ plaque forming units (pfu) were incubated in serum free medium containing either DMSO or ten-fold dilutions of the Thapsigargin, for 1.5 h at 37°C. Thereafter, the samples were chilled at 4°C and diluted by 10^−3^-, 10^−4^- and 10^−5^-fold before being applied onto Vero/MDBK cells in 6-well plates for plaque assaying. The results were plotted as relative infectivity of virions against concentrations of the compound used.

### Virus step-specific assays

Time-of-addition and virus step-specific assays (attachment, entry, RNA synthesis and budding) employed in this study were conducted as previously described (24).

### Overexpression of SERCA

HeLa/goat kidney cells were transfected in 24 well plates, in triplicates with either 1 µg of empty vector (pCR3) or with a plasmid expressing SERCA2 (pCR3-SERCA2) using Lipofectamine 3000 Transfection reagent described above. At 48 h post-transfection, cells were infected with NDV/PPRV at MOI of 1. Virus particles released in the supernatant at 24 hours post-infection (hpi) (NDV) or 96 hpi (PPRV) were quantified by plaque assay.

### Immunofluorescence assay

Vero cells were grown in chamber slides at ∼20% confluency and infected with PPRV/NDV at MOI of 5 for 2 h followed by washing with PBS and replacing with fresh medium. Thapsigargin was applied at 3 hpi and 10 hpi respectively in NDV and PPRV infected cells. The intracellular localization of viral proteins in the virus-infected cells was detected by immunofluorescence assay. Briefly, cells were fixed with 4% paraformaldehyde for 15 min and blocked by 1% bovine serum albumin for 30 min at room temperature. After washing with PBS, cells were stained with primary antibody (rabbit anti-NDV/rabbit anti-PPRV or mouse SERCA2 ATPase) for 30 min in the presence of 0.2% saponin. Thereafter, cells were washed three times with PBS and incubated with a secondary fluorescein isothiocyanate-conjugated anti-rabbit antibody or secondary Rhodamine-conjugated anti-mouse antibody in the presence of 0.2% saponin for 30 min. After being washed again with PBS, the cells were mounted with a medium containing 4,6-diamidino-2-phenylindole (DAPI; Sigma) and examined by fluorescence microscopy.

### Selection of Thapsigargin-resistant viral mutants

PPRV and NDV were sequentially passaged (70 passages) in Vero cells in the presence of either 0.25 µM Thapsigargin or 0.05% DMSO (vehicle control). At each passage, confluent monolayers of Vero cells were infected with the respective virus, washed 5 times with PBS before a fresh aliquot of DMEM was added and incubated for 72-120 h or until the appearance of cytopathic effect (CPE) in ≥75% cells. The virus released in the supernatant was termed as passage 1 (P1). The virus was quantified by plaque assay and it (at MOI of 0.01) was used in the second round of infection, which was termed as passage 2 (P2). Seventy passages of virus infection were similarly carried out. In order to study the relative resistance against Thapsigargin at various passages, Thapsigargin-passaged and DMSO-passaged viruses were used to infect Vero cells at MOI=0.1 and grown in presence of either 0.05% DMSO or 0.5 µM Thapsigargin. The virus released in the supernatant was quantified by plaque assay and fold inhibition levels were determined.

## RESULTS

### SERCA inhibitor blocked paramyxovirus replication

Thapsigargin is a potent inhibitor of SERCA (25, 26). It was identified from a protein kinase inhibitor library described previously (6). In order to determine the *in vitro* efficacy of Thapsigargin against paramyxoviruses (PPRV and NDV) and DNA viruses (BHV-1 and BPXV), we first determined its cytotoxicity in cultured Vero/MDBK cells by MTT assay. As shown in **Fig. 1a** and **1b**, Thapsigargin at concentrations from 0.00064 µM to 2 µM did not significantly affect the cell viability even when incubated in cell cultures for 96 h. However, at higher concentrations (>2 µM), it was found to be toxic to the cells. A non-cytotoxic concentration (0.5 µM) of Thapsigargin was used thereafter.

**Fig. 1.**
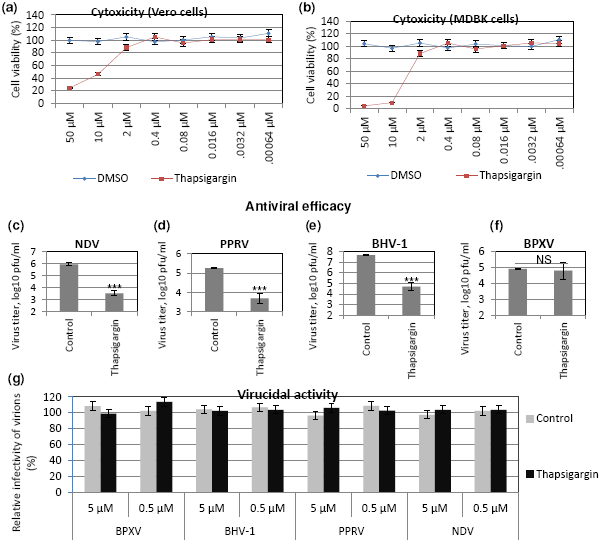
Antiviral efficacy of Thapsigargin. *Cytoxicity (MTT assay):* Five-fold serial dilutions of Thapsigargin (50 µM-0.00064 µM) or equivalent volume of vehicle control (DMSO) were incubated with cultured Vero/MDBK cells for 96 h and percentage of cell viability was measured by MTT assay. Cell viability of Vero (a) and MDBK (b) cells is shown. ***Antiviral efficacy***: Vero/MDBK cells were infected with respective viruses at MOI=0.1 in the presence of 0.5 µM Thapsigargin or vehicle control (DMSO). Progeny virus particles released in the supernatants were quantified by plaque assay. Antiviral efficacy of Thapsigargin against NDV (c), PPRV (d), BHV-1 (e) and BPXV (f) is shown. ***Virucidal activity***: Ten-fold dilutions of the Thapsigargin or DMSO were mixed with ∼10^6^ pfu of the indicated viruses and incubated for 1.5 h at 37°C. Thereafter, the residual viral infectivity was determined by plaque assay in Vero (NDV, PPRV, and BPXV) or MDBK (BHV-1) cells (g). Error bars indicate SD. Pair-wise statistical comparisons were performed using Student’s t test (*** = P<0.001). NS represents no statistical significance.

In order to determine the *in vitro* antiviral efficacy of Thapsigargin, we measured the yield of infectious PPRV/NDV/BHV-1/BPXV in the presence of 0.5 µM inhibitor or vehicle control (DMSO). Thapsigargin significantly inhibited paramyxovirus viz; NDV **(Fig. 1c)**, PPRV **(Fig. 1d)** replication. It also significantly inhibited BHV-1 **(Fig. 1e)** but not BPXV **(Fig. 1f)** replication (DNA viruses), suggesting its potent antiviral activity against paramyxoviruses and BHV-1 virus.

Furthermore, in order to determine whether the antiviral efficacy of Thapsigargin against paramyxoviruses is partially due to direct inactivation of the cell free virions, we incubated the infectious virions with either 0.5 µM or 5 µm Thapsigargin for 1.5 h and subsequently tested the residual infectivity on Vero/MDBK cells. Thapsigargin did not exhibit any virucidal effect on any of the prototype virus tested **(Fig. 1g)** suggesting that the antiviral activity of Thapsigargin is due to the inhibitory effect on virus replication in the target cells.

### SERCA facilitates paramyxovirus replication

In order to further confirm the role of SERCA in virus replication, HeLa/goat kidney cells were transfected with plasmid expressing SERCA2 (pCR3-SERCA2), followed by viral infection. As compared to control plasmid (empty vector)-transfected cells, overexpression of SERCA2 not only facilitated NDV/PPRV replication, but also rescued the inhibitory effect of Thapsigargin on virus replication suggesting that SERCA2 supports paramyxovirus replication **(Fig. 2a** and **2b)**.

**Fig. 2.**
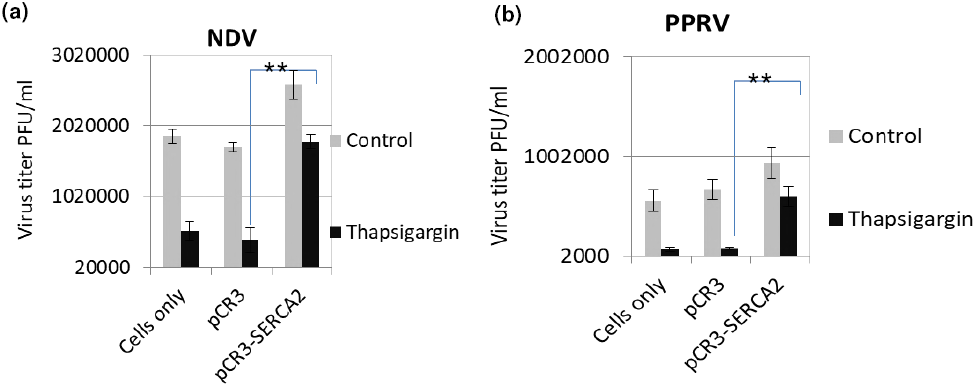
SERCA facilitates paramyxovirus replication. HeLa/goat kidney cells were transfected with SERCA expressing plasmid (pCR3-SERCA2) or control plasmid (pCR3, empty vector). At 48 h-post transfection, cells were infected with NDV/PPRV at MOI of 1. Virus released in the supernatant at 24 hpi (NDV) (a) or 96 hpi (PPRV) (b) was quantified by plaque assay. Error bars indicate SD. Pair-wise statistical comparisons were performed using Student’s t test (** = P<0.01).

### Paramyxoviruses induce SERCA expression

SERCA2 is expressed by most cell types. We also evaluated whether virus infection induces any alteration in SERCA expression. PPRV infection of Vero cells resulted in enhanced SERCA2 expression. As compared to mock-infected cells, a significant induction in SERCA2 expression was observed at 3 hpi, which remained at the peak level between 24-72 hpi, before started declining at 96 hpi **(Fig. 3a, upper panel).** However, the levels of house keeping control gene (GAPDH) were similar at all the time points, suggesting that the enhanced levels of SERCA2 expression were related to viral infection **(Fig. 3a, lower panel).** Besides, we also observed that virus-induced SERCA2 expression could be blocked by Thapsigargin treatment **(Fig. 3b)**.

**Fig. 3.**
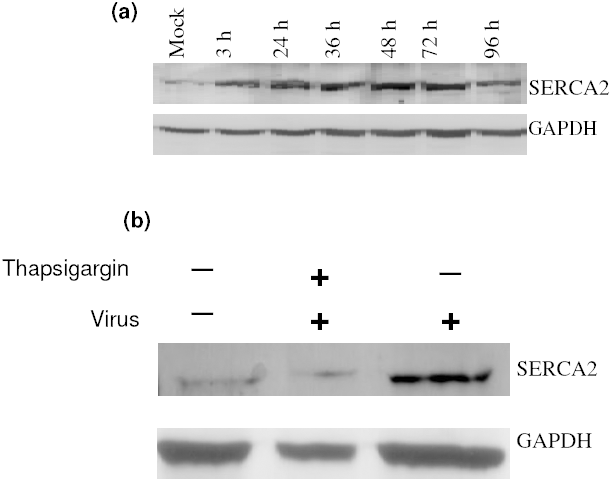
Paramyxovirus infections induce SERCA expression. *(a)* ***PPRV infection induces SERCA2 expression:*** Vero cells were infected with PPRV at MOI 5 and the cell lysates were prepared at indicated time points. The levels of SERCA2 (upper panel) or house-keeping control protein (GAPDH) (lower panel) were examined by Western blot analysis. ***(b) Thapsigargin inhibits NDV-induced SERCA2 expression in Vero cells:*** Vero cells were infected with NDV at MOI 5 for 1 h followed by washing with PBS and addition of fresh medium containing Thapsigargin or vehicle control. Cell lysates were prepared at 16 hpi to probe SERCA2 and GAPDH by Western blot analysis.

### Time-of-addition assay

In order to ascertain the stage(s) of the viral life cycle which can be impaired by Thapsigargin, we performed a time-of-addition assay (one-step growth curve), in which the inhibitor was applied at different time post-infection and the virus released into the supernatant was quantified by plaque assay. The NDV and PPRV varies in their length of replication cycle, ∼10 h and ∼24 h respectively, therefore time-of-addition of inhibitor and time of virus harvest varied from virus to virus. As shown in **Fig. 4a,** the magnitude of viral (NDV) inhibition gradually decreased. The highest inhibition was observed when the inhibitor was applied 30 min before infection. The inhibition levels progressively decreased from 1 hpi to 6 hpi. Thapsigargin did not exhibit any inhibitory effect on virus replication if it was applied at 10 hpi, a later time point in NDV life cycle when the virus is presumably undergoing budding. Similar findings were observed with PPRV; highest inhibition in viral titers was observed when the inhibitor was applied 30 min prior to infection, magnitude of inhibition progressive decreased from 4 hpi to 24 hpi **(Fig. 4b).** The time-of-addition assay, therefore suggested that Thapsigargin may inhibit multiple pre-budding steps of paramyxovirus replication.

**Fig. 4.**
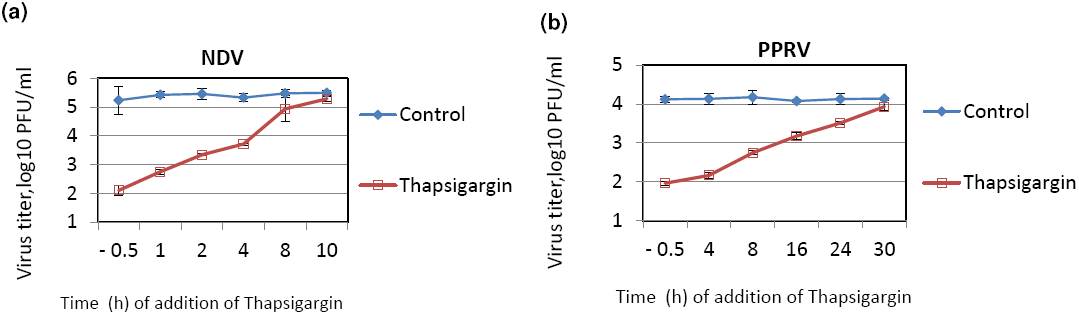
Time-of-addition assay. Confluent monolayers of Vero cells were infected, in triplicates, with PPRV/NDV at MOI of 5, washed 6 times with PBS and fresh medium having either 0.5 µM Thapsigargin or 0.05% DMSO were added at indicated times. Supernatants were collected at 12 hpi (NDV) or 48 hpi (PPRV) and quantified by plaque assay. Time-of-addition assay for NDV (a) and PPRV (b) is shown.

### Thapsigargin does not affect virus attachment, RNA synthesis and budding

Virus step-specific assays viz; attachment, entry, RNA synthesis, subcellular localization of viral proteins and budding were conducted as per the previously described methods (24). Thapsigargin was not found to affect virus attachment, RNA synthesis and budding (data not shown).

### SERCA inhibition impedes paramyxovirus entry

For viral entry assay, pre-attached virus (4°C) was allowed to enter at 37°C in presence of Thapsigargin and infectious virus released in the cell culture supernatant was measured. Thapsigargin treatment resulted in reduced NDV (**Fig. 5a**) and PPRV (**Fig. 5b**) titers suggesting that SERCA inhibitor blocks paramyxovirus entry.

**Fig. 5.**
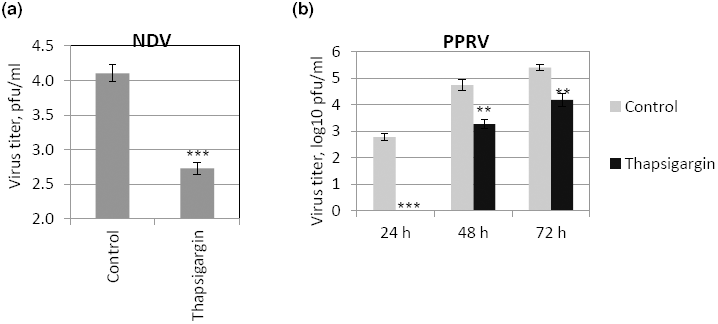
Viral entry. Vero cell monolayers were pre-chilled to 4°C and infected with the respective viruses at MOI of 5 in Thapsigargin-free medium for 1h at 4°C to permit attachment, followed by washing and addition of fresh medium containing Thapsigargin or vehicle-control. Entry was allowed to proceed at 37°C for 1h after which the cells were washed again with PBS to remove any extracellular viruses and incubated with cell culture medium without any inhibitor. The progeny virus particles released in the cell culture supernatants in the treated and untreated cells were titrated by plaque assay. Error bars indicate SD. Pair-wise statistical comparisons were performed using Student’s t test (** = P<0.01,*** = P<0.001).

### SERCA inhibition results in defective subcellular localization of viral proteins

To further examine whether SERCA inhibitor may impact other intermediate step(s) of viral replication, immunofluorescence assay was performed to monitor the subcellular localization of PPRV/NDV proteins in the cytoplasm of the infected cells. The inhibitor or respective vehicle control were applied at 3 hpi (NDV) or 10 hpi (PPRV); a time point when the early events of virus replications (attachment, entry and RNA synthesis) are believed to occur. At 12 hpi (NDV) or 36 hpi (PPRV), when the progeny virus particles are supposed to accumulate at plasma membrane for budding, we observed cytoplasmic foci in majority of Thapsigargin-treated cells **(Fig. 6a** and **6b**). Contrary, in majority of the DMSO-treated cells, virions were found to be localized at the cell surface **(Fig. 6a** and **6b).** Further, the cells with normal (cell surface) or defective (cytoplasmic foci) localization were quantified. Thapsigargin treatment showed defective localization in ∼70% of the cells, as compared to DMSO control wherein this proportion was 10-30 % **(Fig. 6c** and **6d),** suggesting that the SERCA is required for proper localization of the viral proteins (cytoplasm to the cell surface) in infected cells.

**Fig. 6.**
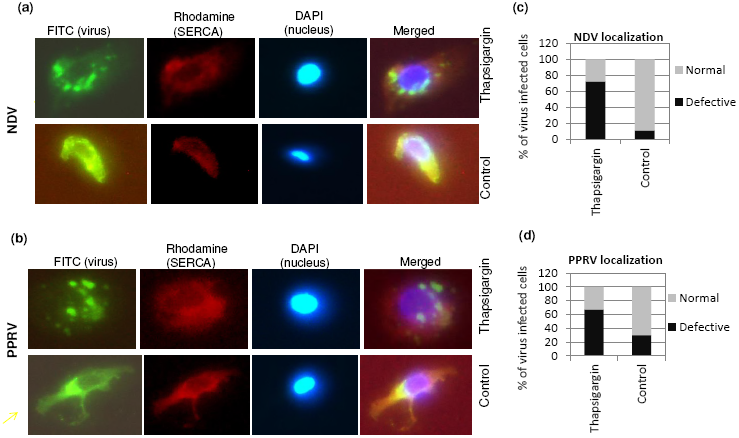
Thapsigargin affects subcellular localization of viral proteins in cytoplasm. Vero cells at ∼ 20% confluence were infected with NDV/PPRV at MOI of 10. At 3 hpi (NDV) or 10 hpi (PPRV) cells were incubated with either 0.5 µM Thapsiargin or 0.05% DMSO. At 10 hpi (NDV) or 36 hpi (PPRV), cells were fixed and subjected for immunofluorescence assay for localization of the viral proteins in the virus-infected cells. Virus was stained with FITC (green) whereas SERCA was stained with Rhodamine conjugate. DAPI was used as nuclear stain. Subcellular localization of NDV (a) and PPRV (b) is shown. Virus infected cells (FITC) were usually of two types-with or without punctuate structures. To quantitate the subcellular localization, 100 cells (under several different fields) were randomly counted. The percentages of the cells with (defective subcellular localization) or without (normal) punctuate structures in Thapsigargin-treated and control-treated is shown for NDV (c) and PPRV (d).

### Selection of Thapsigargin-resistant viral mutants

Due to the high genetic barrier to resistance, host-targeting agents provide an interesting perspective for novel antiviral strategies, rather than the directly-acting agents. NDV, when passaged sequentially in presence of a SERCA inhibitor (Thapsigargin, a host-targeting agent) did not generate a completely resistant phenotype against Thapsigargin, even upon 70 passages in Vero cells (**Fig. 7a**). However, resistance began appearing at ∼P25 and significant resistance was observed at P35 (∼100 fold-inhibition compared to ∼10,000 fold inhibition at zero passage) after which it became stable without acquiring complete resistance (**Fig. 7a**). Fluctuations in the overall viral titers and hence variation in fold inhibition was observed (**Fig. 7a),** which might be due to the fact that the harvest time (72-120 h) and MOI (0.01-0.001) used was not similar at each passage. Therefore, we carried out an experiment wherein a similar MOI (MOI=0.1) and harvest time (24 h) was used to evaluate the relative resistance in Thapsigargin-passaged and control-passaged viruses. As compared to P0 and P70-Control viruses, P70-Thapsigargin virus exhibited significantly lower sensitivity to Thapsigargin, though a completely resistant phenotype could not be observed (**Fig. 7b).** However, all PPRVs (P0, P70-Thapsigargin and P70-Control) revealed similar sensitivity to Thapsigargin even at passage level 70 **(Fig. 7c),** suggesting that PPRV is unlikely to develop Thapsigargin resistant mutants upon long-term passage. Control-passaged viruses did not exhibit any significant resistance against Thapsigargin even upon 70 passages **(Fig. 7b** and **7c)** suggesting that resistance against Thapsigargin (NDV) is not a general phenomenon due to sequential high passages but rather a specific event acquired in presence of Thapsigargin.

**Fig. 7.**
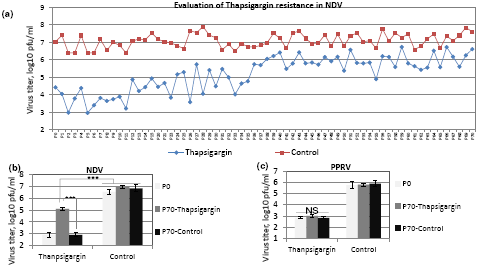
Selection of Thapsigargin-resistant viral mutants. Vero cells were infected with NDV/PPRV at MOI of 0.01 (except at few occasion when it was 0.001 due to extremely low virus yield in cell cultures with Thapsigargin treatment) and grown in presence of either 0.25 µM of Thapsigargin or 0.05% DMSO (vehicle control). The progeny virus particles released in the supernatant were harvested either at 72-120 hpi or when ∼75% cells exhibited CPE. Seventy (70) such sequential passages were made. **(a)** The levels of NDV inhibition at various passage levels in Thapsigargin-treated and untreated cells are shown. **(b) S*ensitivity of long-term Thapsigarin-passaged NDV to Thapsigagrin:*** Vero cells, in triplicate, were infected with P0, P70-Thapsigarin or P70-Control passaged viruses at MOI of 0.1 in presence of either 0.5 µM Thapsiargin or 0.05% DMSO and the progeny virus particles released in the supernatant at 24 hpi were quantified **(c) S*ensitivity of long-term Thapsigarin-passaged PPRV to Thapsigagrin:*** Vero cells, in triplicate, were infected with P0, P70-Thapsigarin or P70-Control passaged viruses at MOI of 0.1 in presence of either 0.5 µM Thapsiargin or 0.05% DMSO and the progeny virus particles released in the supernatant were quantified by plaque assay. Error bars indicate SD. Pair-wise statistical comparisons were performed using Student’s t test (*** = P<0.001). NS represents no statistical significance.

## DISCUSSION

High mutation rate in RNA viruses induces resistance to antiviral drugs and preexisting immunity. The rise in incidence of drug resistance has prompted a shift towards the development of novel antiviral drugs. As compared to the viral genome, genetic variability of the host is quite low and therefore host-targeting agents are considered to impose a higher genetic barrier to generation of resistant viruses (24, 27-29). Thus, a potentially better approach for development of novel antiviral therapeutics would be to target host factors required for viral replication. Targeting host factors could have a significant impact on multiple virus genotypes (strain/serotype) and provide broad spectrum inhibition against different families of viruses which might use the same cellular pathway(s) for replication (24, 30-33). This novel approach has led to the development of some promising compounds for treatment of HCV and HIV (34, 35). In this study, we have shown that targeting SERCA (a Ca^2+^ ATPase) by a small molecule chemical inhibitor-Thapsigargin can block paramyxovirus replication at the level of viral entry and localization of viral proteins in infected cell. We were able to verify the specific requirement of SERCA in paramyxovirus replication; overexpression of SERCA2 rescued the inhibitory effect of Thapsigargin. Therefore, SERCA may present as a novel target for antiviral drug development. When we were writing this manuscript, Hoffmann and coworkers identified that SPCA1 [a secretary pathway calcium (Ca^2+^) transporter that facilitates Ca^+2^ and Mn^2+^ uptake into the trans-Golgi network] also facilitates replication of the members of the family *Flaviviridae, Togaviridae* and *Paramyxoviridae* (21) suggesting that multiple calcium transporters may involved in efficient replication of the paramyxoviruses.

It is generally believed that viruses do not acquire resistance against host-targeting antiviral agents (1, 31, 36, 37). However, in a recent study (38), Schaar and colleagues identified Coxsackievirus B3 (CVB3) mutants that replicate efficiently in the presence of several potent antiviral drugs known to inhibit phosphatidylinositol-4-kinase IIIα (PI4KIIIα), a key cellular factor for CVB3 replication. The authors observed that a single point mutation in the viral 3A protein confers resistance and the drug resistant escape mutants of CVB3 can replicate in cells with low PI4KIIIα. Additionally, cyclosporine A (CsA) resistant hepatitis C virus (HCV) mutant has also been identified (39, 40). In our study, resistance acquired by NDV against SERCA inhibitor adds another example to a short list of viruses which can acquire resistance to host-targeting antiviral agents. To the best of our knowledge, this is the first documented example where a paramyxovirus significantly bypassing its dependency on a cellular factor that is targeted by a small molecule inhibitor. While not yet understood, one possible mechanism underlying acquisition of drug resistance is due to change in host factor requirement (41). For example, under selection pressure in CLDN1 (tight junction protein claudin-1 which serves as an entry factor for HCV)-knock-out cells, CLDN1-dependent HCV evolved to use alternate host factors viz; CLDN6 or CLDN9 (41). Alternatively, resistant viruses may simply have enhanced affinity for its natural substrate, thereby allowing the virus to propagate despite reduction in concentration of the cellular factors (42). Though a complete Thapsigargin resistant NDV phenotype could not be achieved even up to passage level 70, it is a matter of further study how NDV became less dependent on SERCA (under the selection pressure of Thapsigargin). Additional studies are also required to determine which viral proteins(s) have acquired mutations upon prolonged passage in presence of SERCA inhibitor and how these mutations confer resistance against Thapsigargin. In immunofluorescence assay, we could not observe a perfect co-localization of SERCA and viral proteins. The co-immunoprecipitation assay to analyze the interaction between SERCA/virus was unsuccessful, therefore, in this study, we could not determine any direct interaction between SERCA and viral proteins.

To conclude, we have provided strong evidence for SERCA as a crucial host factor in facilitating optimal paramyxoviral replication, thus validating this as a candidate drug target for the development of antiviral therapeutics.

## ACKNOWLEDGEMENTS

This work was supported by the Science and Engineering Research Board, Department of Science and Technology, Government of India [Grant number SB/SO/AS-20/2014].

